# Structure of recombinant formate dehydrogenase from *Methylobacterium extorquens* (MeFDH1)

**DOI:** 10.1101/2023.08.02.551734

**Authors:** Junsun Park, Yoonyoung Heo, Byoung Wook Jeon, Mingyu Jung, Yong Hwan Kim, Hyung Ho Lee, Soung-Hun Roh

**Author notes:** Correspondence to; (H.H.L.); (Y.H.K.); (S.-H.R.). These authors contributed equally to this work (Junsun Park, Yoonyoung Heo, Byoung Wook Jeon).

## Abstract

Formate dehydrogenase (FDH) is critical for the conversion between formate and carbon dioxide. Despite its importance, the structural complexity of FDH and difficulties in the production of the enzyme have made it difficult to elucidate its unique physicochemical properties. Here, we purified recombinant *Methylobacterium extorquens* AM1 FDH (MeFDH1) and used cryo-electron microscopy to determine its structure. We resolved a heterodimeric MeFDH1 structure at a resolution of 2.8 Å, showing a noncanonical active site and a well-embedded Fe-S redox chain relay. In particular, the tungsten bis-molybdopterin guanine dinucleotide active site showed an open configuration with the flexible C-terminal cap domain, suggesting structural and dynamic heterogeneity in the enzyme.

**SIGNIFICANCE:**

- A recombinant MeFDH1 from an inducible expression system
- Structural characterization of recombinant MeFDH1 for its catalytic activity
- A dynamic, open configuration of the C-terminal cap domain

## INTRODUCTION

Formate dehydrogenase (FDH) oxidizes formate to carbon dioxide with the transfer of electrons to an electron acceptor^1-3^. This enzyme, which is important for maintaining cellular redox balance and energy metabolism, is conserved widely from bacteria to many eukaryotes^3,4^. When alternative electron acceptors are present, FDH is essential for microbial metabolism in aerobic environments (e.g., *Methylobacterium extorquens* FDHs used for methanol assimilation) or anaerobic environments (e.g., *Escherichia coli* FDHs used for nitrogen metabolism, and *Shwanella oneidensis* FDHs with fumarate reductase activity). By allowing microorganisms to efficiently use formate as an electron donor, FDH supports their survival and adaptation in diverse ecological niches^5^.

Although FDH enzymes vary across species, they typically consist of multiple subunits that each contribute to functionality. The active site of FDH contains a cofactor, often a metal ion such as molybdenum or tungsten, that catalyzes the oxidation reaction^6^. FDH has applications in bioenergetics^7,8^, biotechnology^9,10^, and environmental science^11^, particularly in the industrial production of formate-based chemicals. Further, researchers are testing FDH for the development of redox enzyme-based biofuel cells^12^. Understanding the structure, function, and mechanism of action of FDH provides valuable insights into the principles of enzymatic catalysis for both industrial and research applications. The structural biology and bioelectrochemistry of *Rhodobacter capsulatus* FDH (RcFDH, PDB ID: 6TGA)^13^ and the native *M. extorquens* AM1 (MeFDH1, PDB ID: 7VW6)^14^ have shown distinct assemblies and electron transfer pathway chains providing mechanistic insights into the enzymatic reactions. However, the structural characteristics of various FDHs and the dynamics of their function are unclear, thereby limiting their applications.

Ongoing research to identify new applications for FDHs in various biotechnological and environmental contexts advances our understanding of this essential biological process. Here, we characterized the structure of a recombinant tungsten-containing FDH from *M. extorquens* AM1 (MeFDH1). We identified and characterized the original strain^15,16^, engineered it with a protein overexpression system, and characterized the recombinant MeFDH1 by single-particle cryo-EM to resolve the 2.8 Å near-atomic structures. We identified an electro-relay system that is highly conserved in this enzyme. In addition, we found structural flexibility in the C-terminal cap domain, which can accommodate the tungstate bis-molybdopterin guanine dinucleotide (W-bis-MGD) ligand in the active site ^17^.

## RESULTS

### Generation of recombinant MeFDH1

Previously, Yoshikawa et al., ^14^ determined the structure of the native FDH complex from *M. extorquens* AM1 (MeFDH1). Here, we overexpressed a recombinant FDH incorporating a His tag for purification. Because we could not produce MeFDH1 using an *E. coli* expression system, we used *M. extorquens* AM1 for protein overexpression. We knocked out the endogenous *fdh1a* and *fdh1b* genes in *M. extorquens* AM1 and introduced the methanol-inducible pCM110-*fdh1a/b*-His plasmid into the strain (Fig. 1A). The expressed protein was purified by a combination of affinity, ionic exchange, and size exclusion chromatography (SEC). We used SEC and multi-angle light scattering (SEC-MALS) experiments to show that MeFDH1 is a heterodimer of alpha and beta subunits (∼169 kDa). Furthermore, sodium dodecyl sulfate-polyacrylamide gel electrophoresis (SDS-PAGE) analysis confirmed the presence of alpha and beta proteins, indicating the success of the recombinant system in producing a stable complex (Fig. 1A).

**Fig. 1.**
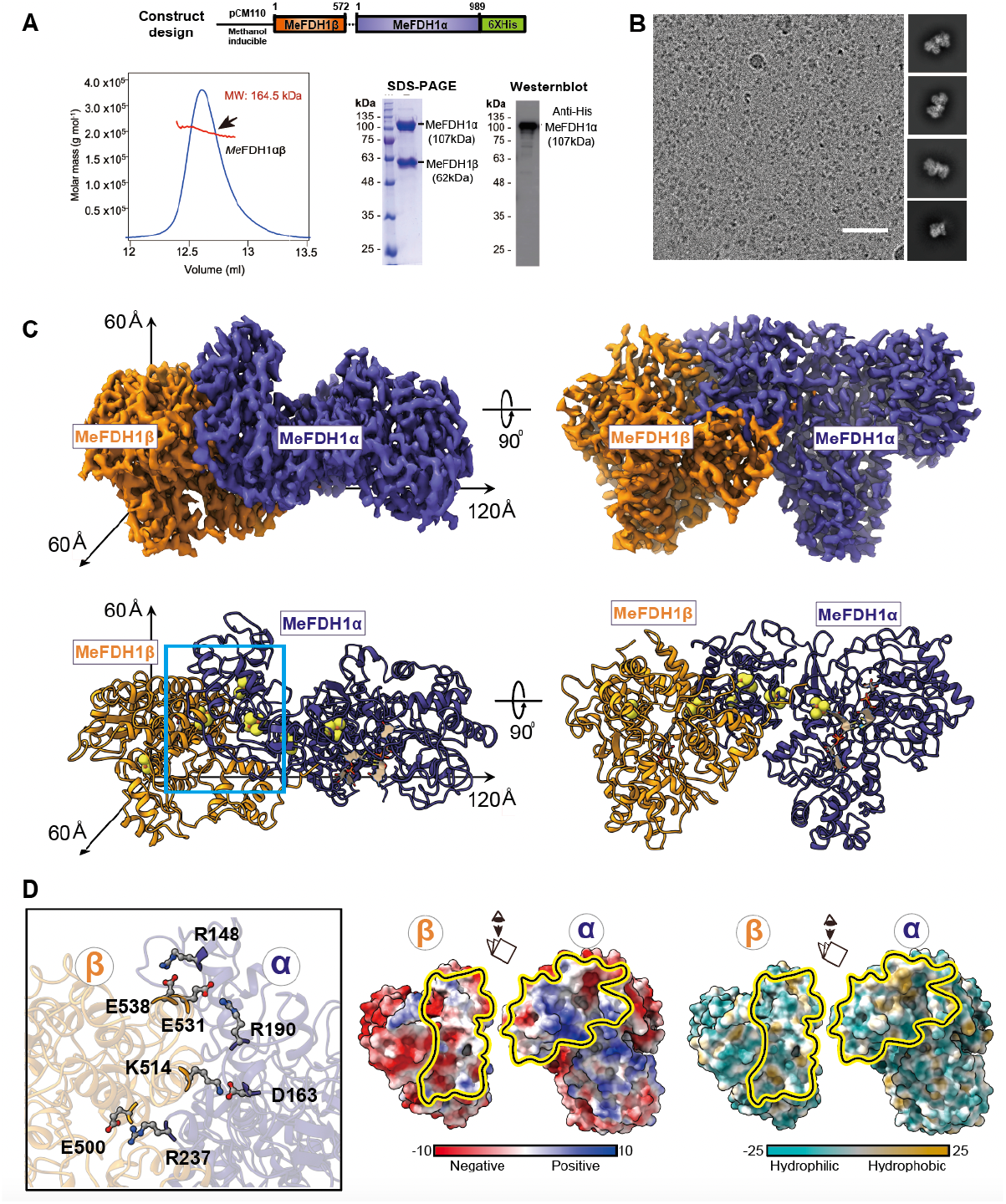
Cryo-EM structure of the recombinant MeFDH1 complex. (A) Plasmid design for expression of recombinant MeFDH1 (top). SEC-MALS data for the oligomeric state of MeFDH1. The thick line represents the measured molecular mass (164.5 kDa). SDS-PAGE of purified MeFDH1 and western blotting of purified MeFDH1 oligomers using an anti-His antibody. (B) A representative cryo-EM micrograph and 2D averages of MeFDH1. The scale bar = 50 nm. (C) 3D reconstruction of the MeFDH1 binary complex and the corresponding atomic model. The dimensions along each axis are shown by the arrows. (D) The interface analysis of MeFDH1-α and MeFDH1-β. Interacting residues are shown as balls and sticks (left). Electrostatic and hydrophobic surface analysis of the interface (right). Yellow lines indicate the contact surfaces between the two subunits.

### The overall architecture of recombinant MeFDH1

We performed single-particle cryo-EM to visualize the molecular architecture of the recombinant MeFDH1. Based on 640,000 particles, the 2D class averages of particle images showed that they were well converged for the particle populations (Fig. 1B and S1A). After multiple rounds of 3D particle classification, we successfully obtained two converged maps of holoenzyme MeFDH1, yielding an overall resolution of 2.8 Å. In addition, we obtained a partial map at 3.17 Å that has beta and part of the alpha protein (residues 1∼287) (Fig. S1B). The consensus holoenzyme map displayed two distinct and isolated densities with clearly visible side chains for most of the protein area (Fig. S1E). We fitted and refined the atomic models based on side chain densities using MeFDH1-α/β complex homology model from the Swiss Model^18^.

Our structure showed a heterodimeric architecture of MeFDH1-α and MeFDH1-β subunits with overall dimensions of 60 × 60 × 120 Å (Fig. 1C). The alpha subunit comprises the active site and the electron transfer funnel shielded by the outer secondary structures. The beta subunit comprises the diaphorase unit and the electron transfer funnel, in which two subunits are arranged in a row in the dimer. MeFDH1-α and MeFDH1-β share a large surface contact over ∼2000 Å^2^ as determined using the PISA program^19^. The residues at the interface comprise about 50% of the polar residues, including multiple complementary intermolecular salt bridges (αR148-βD538, αD163-βK514, αR190-βE531, and αR237-βE500, Fig. 1D). Notably, these charged residues are well conserved throughout FDHs (Fig. S2A). Our results suggest that electrostatic complementarity stabilizes intermolecular contacts and the formation of an effective electron transfer relay for FDHs.

### Electron transfer relay in MeFDH1

FDH catalyzes redox reactions that oxidize formate to carbon dioxide through electron transfer^20^. The MeFDH1-α subunit is essential for catalysis of the substrate, formate. Our structure showed that MeFDH1-α includes W-bis-MGD, three [4Fe-4S] clusters (A1-A3), and one [2Fe-2S] cluster (A4) (Fig. 2A). Each Fe-S cluster was uniquely coordinated by four neighboring cysteine residues. Specifically, two [4Fe-4S] (A2 and A3) and one [2Fe-2S] (A4) were coordinated in the N-terminal domain of the alpha subunit (residues 1−289). The signal intensity for [2Fe-2S] vs. [4Fe-4S] was experimentally distinguishable, and [4Fe-4S] showed ∼50% more significance at the same contour level (Fig. S1E). The MeFDH1-β subunit has an FMN binding domain and shows an overall globular structure containing the ubiquitous diaphorase unit. The MeFDH1-β subunit contains a flavin mononucleotide (FMN) cofactor, a [4Fe-4S] cluster (B1), and a [2Fe-2S] cluster (B2) (Fig. 2A). FMN is coordinated at the center of the MeFDH1-β subunit, and the B1 and B2 clusters are located around the FMN.

**Fig. 2.**
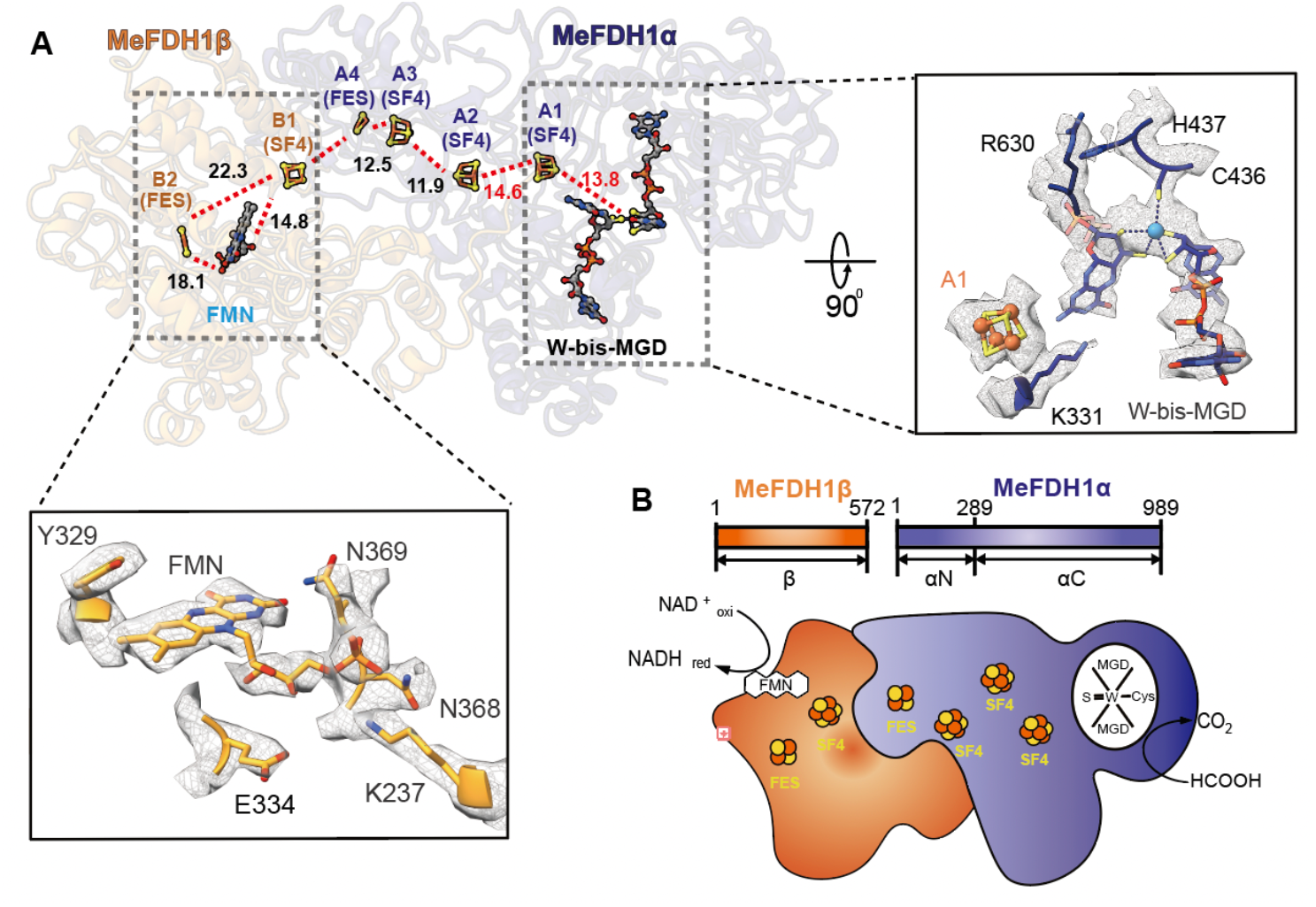
Ligands and the electron transfer relay of MeFDH1. (A) Geometrical arrangement of electronically coupled cofactors (left). Cofactors are shown in the stick model. The distances between the cofactors are given in angstroms for center-to-center measurements. The active site and FMN binding site, as well as interacting residues, are shown with fitting of the model to the electron density map. (B) A schematic diagram of formate oxidation by MeFDH1; each cofactor and Fe-S cluster is shown as a cartoon. FES = [2Fe-2S], SF4 = [4Fe-4S]

The arrangements of the two cofactors (W-bis-MGD and FMN) and five Fe-S clusters (A1, A2, A3, A4, and B1) may serve as an electron relay, such that the electrons flow from the tungsten center (W-bis-MGD) to FMN via a chain of Fe-S clusters spaced optimally for formate oxidation (Fig. 2A). All edge-to-edge distances between cofactors are ∼14 Å, the maximum distance for physiological electron transfers^21^. The [2Fe-2S] cluster B2 in MeFDH1-β, which lies outside of the electron relay (> 15 Å), may not be involved in electron transfer. The geometrical arrangement of electrically coupled cofactors and the electron transfer pathway in MeFDH1 provide a structural mechanism by which MeFDH1 catalyzes conversions between CO_2_ and formate. W-bis-MGD, FMN, and the five Fe-S clusters (A1, A2, A3, A4, and B1) form the redox chain for an electron relay, as shown in other species^13,22^ (Fig. 2B). The W-bis-MGD cofactor, which is an active site for formate oxidation, is surrounded by amino acids that are highly conserved in FDHs (Fig. S2A). The tungsten atom is coordinated by the two dithiolene groups of the bis-MGD molecule as well as the sulfur of Cys436.

### Structural conservation in the FDH family

To determine the evolutionary relationship of MeFDH1 to the FDH family, we compared our structure with others from the DALI database^23^. The N-terminal MeFDH1-α subunit (residue 1−289) is very similar to the HoxU subunit of the NAD^+^-reducing [NiFe] hydrogenase from *Hydrogenophilus thermoluteolus* (PDB ID: 5XF9)^24^. The C-terminus shows high homology with FDH H (FdhF) of *E. coli* (PDB ID: 1FDO)^25^ (Fig. S3A). In particular, *ConSurf* analysis showed that the conserved active site in MeFDH1 coordinated with the metal and the pyranopterin cofactor (Fig. 3A), where the catalytic reaction occurs. MeFDH1-β is very similar to the Nqo1/2 respiratory complex I from *Thermus thermophilus* (PDB ID: 2FUG)^22^, and the HoxF subunit of the [NiFe] hydrogenase from *H. thermoluteolus* and is predicted to be an FMN-bound enzyme for the oxidation of NADH (Fig. S3A). This similarity indicates that MeFDH1-β is an NADH dehydrogenase. In addition, the residues (F232, K237, K351, and E334) in the binding site coordinating NADH are highly conserved among other FDH families (Fig. S4D and E). This conservation suggests that the ligand binding sites of MeFDH1 share a common architecture in metal binding enzymes. Whereas MeFDH1 forms a dimer, the recently characterized *R. capsulatus* FDH (RcFDH) resolved as a dimer of the FdsABGD hetero-tetramer, comprising several functional modules. MeFDH1-α shows structural similarity with FdsA of RcFDH (RMSD of 1.858 Å for 628 Cα residues), whereas MeFDH1-β is similar to the heterodimer of FdsB and FdsG in RcFDH (FdsB: RMSD of 2.20 Å for 158 Cα residues and FdsG: RMSD of 1.876 Å for 110 Cα residues) (Fig. 3B). The N-terminal domains of FdsB and FdsD do not exist in MeFDH1 (Fig. S3B). The arrangements of two cofactors (W-bis-MGD and FMN) and five Fe-S clusters (A1, A2, A3, A4, and B1) that function as an electron relay match the corresponding cofactors in RcFDH (Fig. 3B). The [2Fe-2S] cluster B2 in MeFDH1-β, which lies outside of the electron transfer pathway, corresponds to the G1 cluster in RcFDH. Although the high structural similarity of the cofactors is reflected in a good match of the positioning of the Fe-S clusters in both MeFDH1 and RcFDH, one major difference is the absence of A5 clusters in MeFDH1 (Fig. 3B). As it has been suggested that the A5 cluster lies at the dimer interface of the FdsABGD heterotetramer and allows electron transfer between the two FdsABGD heterotetramers^13^, we infer that the A5 cluster is not necessary for MeFDH1. The shared structural similarities among the domains suggest that the overall architecture of MeFDH1 is common in the FDH family.

**Fig. 3.**
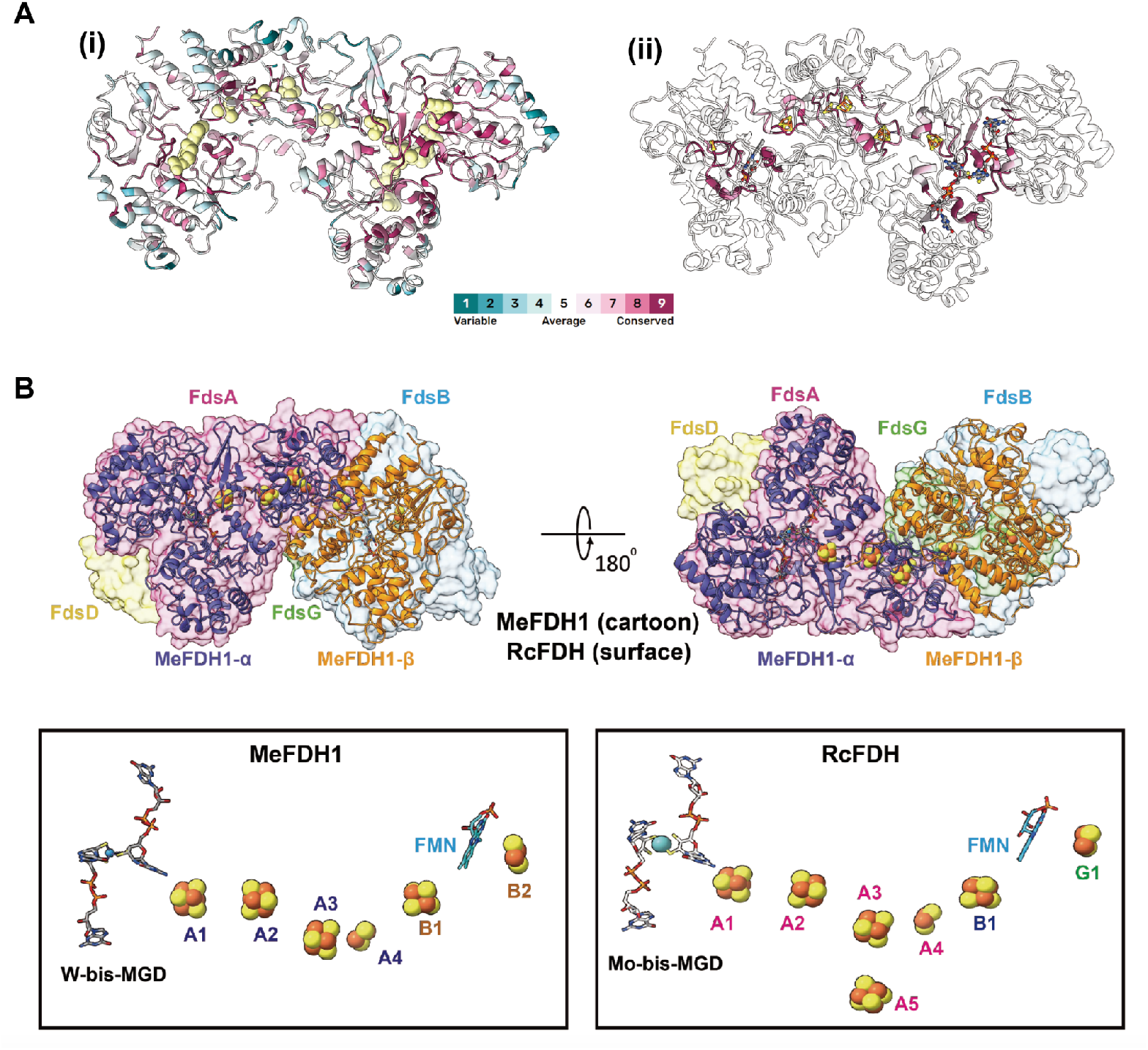
The global cofactor arrangement of MeFDH1 and RcFDH. (A) Conservation score of MeFDH. Left (i) is the overall conservation score, and right (ii) is focused on the region around the FeS clusters. Residues around the FeS clusters are highly conserved. (B) Structural comparison between MeFDH1 and RcFDH. The two structures are aligned, and MeFDH1 is shown as a cartoon, whereas RcFDH is represented as a transparent surface (top). The geometrical arrangement of the electronically coupled cofactors in MeFDH1 and RcFDH. A, B, and G structures indicate Fe-S clusters.

### The dynamic cap domain for the active site

The structures we determined are consistent with the previous report on the structure of MeFDH1^14^; however, we could not visualize the C-terminal cap domain (amino acids 858-989) of the alpha subunit (Fig. 4A). Although we used 3D variability analysis and focused the classification using a mask on the cap domain, we did not observe any notable region of extra electron density. However, SDS-PAGE and western blot analyses confirmed the presence of the C-terminus of the alpha subunit in the complex (Fig. 1A). Therefore, the cap domain is likely highly dynamic, making it challenging to capture its structural details using our approach.

**Fig. 4.**
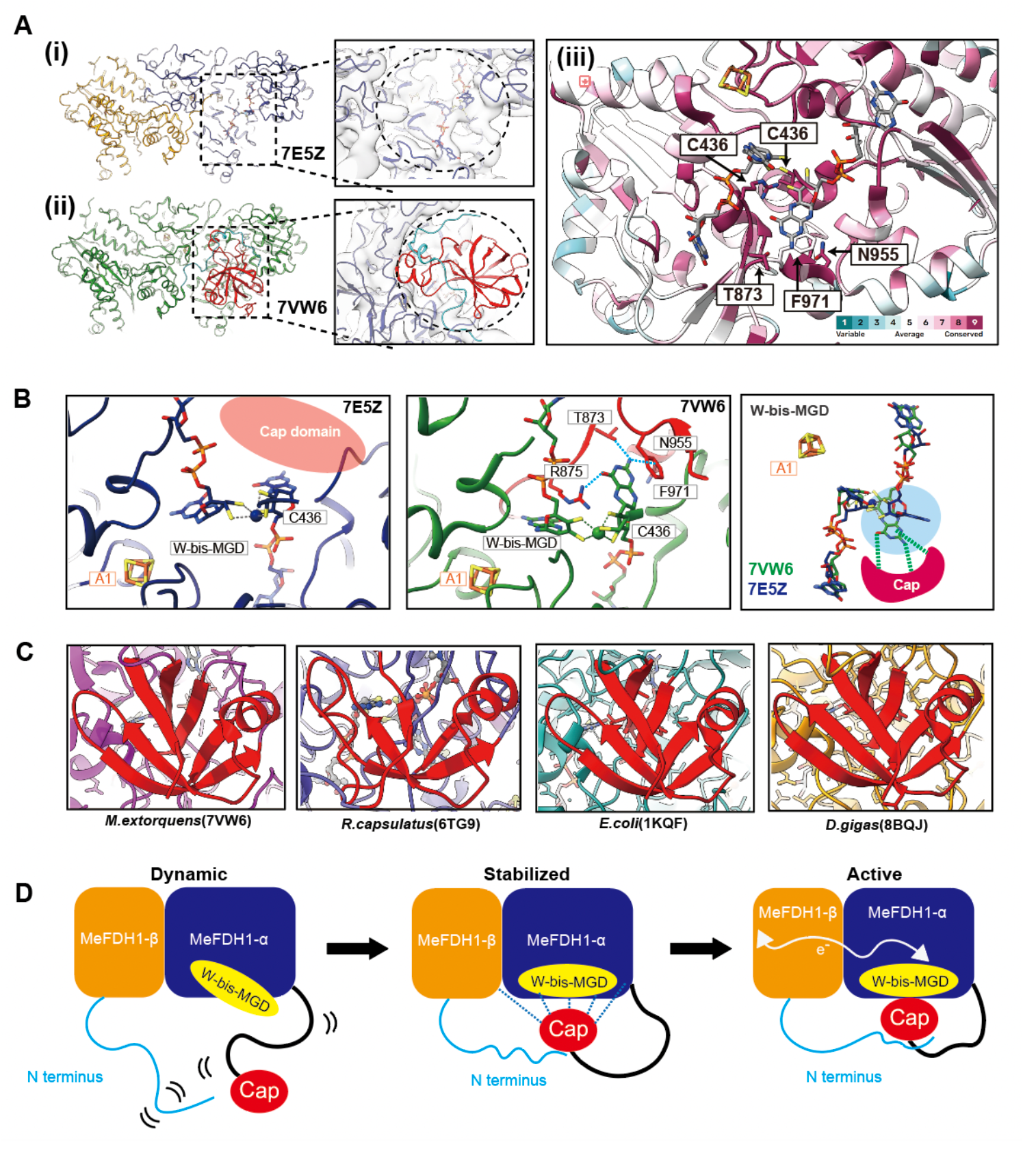
The dynamic cap domain of MeFDH1. (A) MeFDH1 structure models for 7E5Z from this study (i) and 7VW6 (ii) depicted in the same orientation. The dotted box shows the location of the cap domain with a zoomed-in view of the cap domain with an electron density model fitted. (iii) Conservation score of cap domain; residues binding MGD are highly conserved. (B) A zoomed-in view of the active site for each MeFDH1 structure. The cap domain is illustrated by the red schematics. The dotted lines show the interactions between residues and ligands. (C) Structurally conserved cap domains (red) in various species. (D) A schematic model of the dynamic cap domain near the active site and the flexible N-terminus of the β subunit. After W-bis-MGD is loaded to the active site of MeFDH1, the cap domain interacts with the α subunit and the flexible N-terminus of the β subunit. Then the cap domain docks on the active site and stabilizes W-bis-MGD through specific interactions that prevent solvent exposure of the ligand.

To understand the dynamics and functionality of the cap domain, we compared our structure with the previous MeFDH1 structure (PDB: 7VW6)^14^. The 7VW6 structure displays the complete cap domain, which interacts with the N-terminus of the beta subunits, effectively blocking the active site from the solvent (Fig. 4A). However, in our structure, lacking a visible cap domain, the N-terminus of the beta subunits is oriented away from the cap domain and lacks structure (Fig. S4A). Therefore, the active site is exposed to the solvent. Also, the cap domain contributes to the orientation of the active site via certain salt bridges within the alpha subunits and forms salt bridges with K26 and D76 in the N-terminus of the beta subunit, providing further stability to the binding of the cap domain (Fig. S4C). The absence of such interactions in our structure indicates the flexibility of the cap in the absence of these interactions (Fig. S4B).

Our map indicates the dynamic state of the cap domain, which coordinates the protein cofactor^25^. Two structures showed W-bis-MGD density in the active site; however, they showed different MGD binding orientations. Although a pyranopterin near the A1 cluster showed similar binding in two structures, another pyranopterin in 7VW6 was also coordinated by the cap domain through hydrophilic and pi-stacking interactions (Fig. 4B). When two ligands were aligned, the pyranopterins making contact with the cap domain showed significantly different conformations because of the coordination by the cap domain. This suggests the direct stabilization of bis-MGD by the cap domain.

## DISCUSSION

We used cryo-electron microscopy to determine the structure of a recombinant His-tagged FDH from *M. extorquens* AM1, MeFDH1. Our structure resolved a heterodimeric MeFDH1structure at 2.8 Å, showing W-bis-MGD, FMN, and Fe-S clusters forming an electron transfer relay to oxidize formate to CO_2_. In particular, the cap domain near the W-bis-MGD active site showed an open configuration with a flexible C-terminus cap domain, suggesting dynamic and structural heterogeneity.

FDHs show variations in subunit composition, cofactors, and metals across prokaryotes and eukaryotes^6^. MeFDH1 has a distinct heterodimeric structure with a unique array comprising tungsten and an FMN cofactor for electron transfer. However, the conservation scores indicate that the residues surrounding the FeS clusters that form the electron transfer arrays in FDHs are highly conserved (Fig. 3A). The C-terminal residues of the cap domain, which directly stabilizes MGD, are also well conserved. In contrast, there is little sequence-level conservation in the peripheral region of the cap domain (Fig. 4A). Because the residues surrounding the cofactors are highly conserved, it is likely that MeFDH1 shares a catalytic mechanism with other members of the FDH family in the conversions of formate and CO_2_.

Previous work^17^ suggests that dynamic detachment of the cap domain of FDHs from the protein body is required for efficient insertion of the cofactor into the active site during enzyme maturation. The behavior of the cap domain in our study indicates that we observed a transient open configuration of the cap domain following the loading of the cofactor. Next, the cap domain might undergo a conformational change with the closure of the ligand binding site and stabilization of the cap domain by inter- and intra-subunit electrostatic interactions (Fig. 4D). The disordered N-terminus of the beta subunit adopts a rigid structure due to the proximity of the cap domain, and then the cap domain coordinates W-bis-MGD, providing additional stability and shielding the active site from the surrounding solvent. This coordinated interaction results in the formation of the complete electron transfer relay. The high conservation among FDHs of the residues that coordinate bis-MGD in the active site (Fig. 4A) and the cap domain (Fig. 4C) suggests that this mechanism is conserved in similar FDHs. Considering the involvement of specific chaperones in the maturation of FDH^26-28^, the open conformation of MeFDH1 in our study could be attributed to insufficient chaperones for the overexpressed the protein. However, further evidence is required to support this hypothesis.

## ACKNOWLEDGEMENTS

This work was supported by Korea NRF CGRC (2015M3D3A1A01064919), ERC (2020R1A5A1019631), KCRC (2014M1A8A1049296), NRF (2019M3E5D6063871, 2020R1A5A1018081, 2022R1A2B5B02002529, and 2022R1A5A6000760), Creative-Pioneering Researchers Program of Seoul National University and SUHF foundation. Cryo-EM data collected at the Seoul national university cryoEM facilities (Center for Macromolecules and Cell Imaging). We thanks to Dr. Jun Sung-Hoon at KBSI for initial cryoEM data screening.

## AUTHOR CONTRIBUTION

J.-B.W. carried out cloning; H.Y. purified the protein sample; H.Y. and P.J. carried out cryo-EM experiments and P.J. carried out data analysis. P.J. H.Y, J.-B.W, S.-H.R wrote the manuscript. All authors contributed to the final manuscript.

## METHODS

### Generation of recombinant MeFDH1

#### Cloning and cell culture

We used plasmid pCM110 to express recombinant MeFDH1 in a *Δfdh1αβ* mutant of *M. extorquens* AM1, as described previously^1,29^. Plasmid pCM110-MeFDH1 contains the *fdh1b (*GenBank accession ACS42635.1) and *fdh1a* (GenBank accession ACS42636.1) genes encoding MeFDH1, inserted between the methanol-inducible *P*_*mxaF*_ promoter (from the *mxaF*-encoded methanol dehydrogenase) and the T7 terminator, with a His_6_ tag at the C-terminus of the alpha subunit (Fig. 1A). The knockout mutant, lacking *fdh1αβ*, made using a Cre-*lox* knockout system^30^, was used for the expression of MeFDH1 from pCM110-MeFDH1 by growing cells for 20 h at 30 °C to an optical density (OD_600_) of 0.5–0.8 before induction of protein expression with 0.5% methanol. After 48 h, the cells were harvested at 7,000 rpm for 15 min at 4 °C, and cell aliquots were stored at −70 °C until used.

#### Purification of MeFDH1

Frozen *M. extorquens* AM1 cells with pCM110-MeFDH1 were lysed in buffer A (50 mM Tris-HCl, pH 8.0) with 1 mM PMSF by passing them through a microfluidizer and centrifuged at 4,611 × g (Vision V506CA rotor) for 30 min at 4 °C to remove the cell debris. The supernatant was applied to a Ni-NTA affinity column (Qiagen) equilibrated with buffer (50 mM Tris-HCl, pH 8.0, 300 mM NaCl, 10 mM imidazole). The protein was eluted with a linear gradient of 100–500 mM imidazole. Protein fractions were diluted two-fold in buffer A and loaded onto a Source 15Q column (GE Healthcare) equilibrated with buffer A. The protein was eluted with a gradient of 0–600 mM NaCl, and MeFDH1 was eluted at about 300 mM NaCl. Further purification used size exclusion chromatography (HiLoad 16/600 Superdex 200 prep grade, GE Healthcare) on a column that was previously equilibrated with a buffer containing 50 mM Tris-HCl pH 8.0 and 100 mM NaCl. Western blot analysis used anti-His-tag antibodies to confirm the presence of the His tag and the C-terminal cap domain.

#### Size exclusion chromatography with multi-angle light scattering (SEC-MALS)

SEC-MALS experiments for MeFDH1 were performed using an FPLC system (GE Healthcare) connected to a Wyatt MiniDAWN TREOS MALS instrument and a Wyatt Optilab rEX differential refractometer. A Superdex 200 10/300 GL (GE Healthcare) gel-filtration column, pre-equilibrated with buffer A containing 5 mM β-mercaptoethanol, was normalized using ovalbumin. Proteins (1 mg) were injected at a flow rate of 0.4 ml/min. Data were analyzed using the Zimm model for static light-scattering data fitting and graphed using an EASI graph with a UV peak in the ASTRA V software (Wyatt).

### Cryo-EM data collection and image processing

#### Cryo-specimen preparation

We applied 3 μl of purified MeFDH1 (0.5 mg/ml) to a 1.2/1.3 holey carbon 200 mesh grid (Quantifoil), blotted for 4 s in a 100% humidity environment at 12 °C and then froze the specimens by plunging them into liquid ethane using a Vitrobot Mark IV (FEI).

#### Cryo-EM imaging

Collection of image data was performed using a Glacios transmission electron microscope (Thermo Fisher Scientific) operating at an acceleration voltage of 200 kV under parallel illumination conditions. Images were acquired with a Falcon 4 direct electron detector (Thermo Fisher Scientific) at a nominal magnification of 92,000×, corresponding to a calibrated size of 1.08 Å per pixel, with a 50 μm condenser lens aperture. Automated data collection was performed using EPU software (Thermo Fisher Scientific), and 1761 exposures were recorded in a total dose of 52 e^−^/Å^2^, fractionated over 50 movie frames. The detailed imaging conditions are shown in Table S1.

#### Imaging processing

All data processing was performed using cryoSPARC v.3.3.1^31^. Raw movies were imported, and motion was corrected using patch motion correction, and the contrast transfer function (CTF) was estimated using patch CTF estimation, which was implemented in cryoSPARC. A small set of particles was collected by auto-picking without templates and averaged, followed by template picking. Then, intact particles were applied to Topaz training^32^, and 640,408 good particles were extracted with a box size of 240 from the micrographs. Ab initio modeling and 3D heterogeneous refinement yielded four 3D classes representing the MeFDH1 holoenzyme map. The particles of the MeFDH1 holoenzyme were subjected to non-uniform refinement^33^ and then further refined by CTF refinement and local motion correction. The final MeFDH1 holoenzyme map had a resolution of 2.83 Å determined by the gold-standard Fourier shell correlation (FSC) curve at 0.143.

#### Model building and refinement

The initial MeFDH1 map was homology modeled using the Swiss Model server^19^. The map was subjected to density modification in PHENIX *RESOLVE cryo-EM* ^*34*^ to enhance the interpretability of the map. The model was initially fitted into the map in UCSF chimera ^35^ and refined in PHENIX *real-space-refine*, and manually fitted in *Coot* ^36,37^.

## Data availability

The cryo-EM structures have been deposited in the Electron Microscopy Database (EMDB) under accession codes EMD-30995 (https://www.emdataresource.org/EMD-30995) and in the Protein Database (PDB) under accession code 7E5Z (https://www.rcsb.org/structure/7E5Z). Other data are available from the corresponding author upon reasonable request.

